# Cardiovascular parameters in capitive blue-fronted amazon parrots (*Amazona aestiva*, Linnaeus, 1758) with varying body condition scores

**DOI:** 10.1101/712273

**Authors:** Gisele Junqueira dos Santos, Amanda Sarita Cruz Aleixo, Jeana Pereira Da Silva, Alícia Giolo Hippólito, Bárbara Sardela Ferro, Elton Luis Ritir Oliveira, Priscylla Tatiana Chalfun Guimarães Okamoto, Maria Lucia Gomes Lourenço, Vania Maria de Vasconcelos Machado, Alessandra Melchert

## Abstract

This study aims at determining the echocardiographic, radiographic and tomographic parameters of blue-fronted Amazon parrots (*Amazona aestiva*) with varying body condition scores. Thirty-five birds grown in captivity were included in the study and allocated into groups according to their respective body condition scores: Lean, Ideal and Obese. The echocardiographic evaluation revealed that obese parrots presented lower right ventricle dimensions in diastole than lean parrots. The fractional shortening was considerably lower in obese parrots than in parrots with lean and ideal body condition scores, but without statistical significance. The flow rate and the aortic pressure gradient were lower in the lean group than in the ideal group. No differences were observed between the groups when comparing the radiographic and tomographic measurements. Therefore, as is the case in mammals, we can conclude that alterations in the nutritional state of blue-fronted amazon parrots lead to cardiovascular dysfunctions detected only through an echocardiographic evaluation, which represents an important diagnostic tool for these animals. Computed tomography scans allowed a better identification of the structures of the cardiovascular system without the overlaying structures of the celomatic cavity observed in radiographic images. However, radiographic examinations should still be considered the standard screening examination to identify cardiac alterations such as increased or reduced organ dimensions. Standardizing the technique and describing the measurements obtained in this study may serve as a basis for further research.

## Introduction

Blue-fronted Amazon parrots (*Amazona aestiva*, Linnaeus, 1758) are popular pets and are often kept in captivity [1,2], with characteristics such as an attractive appearance, interaction with the owner and a long lifespan helping to explain such popularity [3]. In captivity, it is common for these birds to have diets based on oilseeds, which are rich in fat and low in minerals and vitamins, and may potentially lead to nutritional and cardiovascular disorders, compromising the quality of life of the birds [4,5].

Cardiovascular diseases are common in birds [6], representing a diagnostic challenge [7] because the physical examination may be difficult to interpret correctly [8]. Straub et al. [9] have reported heart injuries in over 30% of the necroscopic examinations conducted on psittacidae. In addition, these diseases may present signs that go undetected by the owners, leading to a dependency on complementary examinations for the diagnosis [10].

It is important to considered that wild birds kept in captivity often have inadequate intakes of vitamins and calcium, with excessive fat intake or low protein and energy intake, which lead to nutritional alterations that may go unnoticed and predispose the animals to cardiac diseases [4]. Heart diseases in pet birds are more significant than previously believed and a cardiac evaluation should be included in the routine clinical examination [11]. The advanced technologies available nowadays, the improvement of anesthetic techniques, the behavioral conditioning and the better understanding of the physiology of these animals have resulted in a fast expansion of the imaging diagnosis methods available to wild animals [12].

Considering that parrots are often taken as pets and receive inadequate diets, which represents a risk factor for cardiovascular diseases, this study aims at describing the echocardiographic, radiographic and tomographic cardiovascular parameters in captive blue-fronted Amazon parrots (*A. aestiva*) with varying body condition scores. As far as the authors know, there are no studies describing the influence of body condition score on the echocardiographic, radiographic and tomographic examinations for this species.

## Materials and Methods

The study was approved by the Ethics Committee on Animal Use (CEUA, *Comitê de Ética em Uso Animal*) at FMVZ, UNESP, Botucatu, Brazil, under protocol number 183/2015 and also by the Chico Mendes Institute of Biodiversity Conservation (ICMBio, *Instituto Chico Mendes de Conservação da Biodiversidade*), by the Biodiversity Authorization and Information System (SISBIO, *Sistema de Autorização e Informação em Biodiversidade*), under protocol number 81121252. A total of 35 adult blue-fronted Amazon parrots (*A. aestiva*) without sex distinction were included in the study, all of which were kept in captivity for at least two years at the Wildlife Medicine and Research Center (*CEMPAS*, *Centro de Medicina e Pesquisa em Animais Selvagens*) at the School of Veterinary Medicine and Animal Science, UNESP, Botucatu-SP, Brazil. The study was conducted from September 2017 to December 2017 and the birds came from seizures conducted by the Brazilian Environmental Police and donations, which means there are no prior history of the health state of these animals. The animals were kept in a room of approximately 40 m² with access to indoors and outdoors areas.

A score system with five tiers was used to establish the body condition scores (BCS) based on the pectoral musculature [13,14]: Score 1: animals with concave musculature over the keel apex, representing malnutrition and thinness; Score 2: moderate muscle mass with the keel apex easily palpable, corresponding to mild thinness; Score 3: attributed to parrots with convex pectoral muscles; Score 4: muscle mass level with the keel apex (scores 3 and 4 are considered ideal); score 5: muscle extend beyond the keel, representing obesity. The accumulation of subcutaneous fat may be felt over the pectoral area, sternum and abdomen, indicating obesity [13]. The birds had their weights measured on digital scales and their lengths measured from the occipital bone until the tip of the tail.

After determining the BCS, each bird was allocated into three distinct groups: Lean Group (n=11), BCS below the ideal for the species (scores 1 and 2); Ideal Group (n=14), ideal BCS for the species (scores 3 and 4); Obese Group (n=10), BCS above the ideal for the species (score 5). During the experiment, the birds received drinkable water and their usual diets (bananas, apples, papayas and pelletized feed for psittacidae, Psita Sticks, Alcon^®^). The parrots did not eat or drink for four hours before the examinations [15] and were anesthetized with intravenous midazolam (Dormonid^®^) 1 mg/Kg, and ketamine (Dopalen^®^) 20 mg/Kg through the jugular vein.

The echocardiographic examination was conducted with a SonoSite device (European Heardquarters, United Kingdom), model M Turbo, with a sectorial transducer, frequency of 7.5 MHz, B-Mode, and Spectral and Color Doppler functions. The examination was conducted with electrocardiographic monitoring, allowing a precise determination of the cardiac cycle. The animals were kept in erect position with a suitable device (Fig 1A) and a four-hour fast was adopted to improve the accuracy of the images since the gastrointestinal tract may compromise the penetration of ultrasound waves [16]. The examination was conducted through the ventromedial approach, with the transducer positioned medially, directly behind the sternum (Fig 1B and 1C). The measurements, including flow rate (m/sec) and aortic pressure gradient (mmHg) were taken through this window.

**Fig 1:**
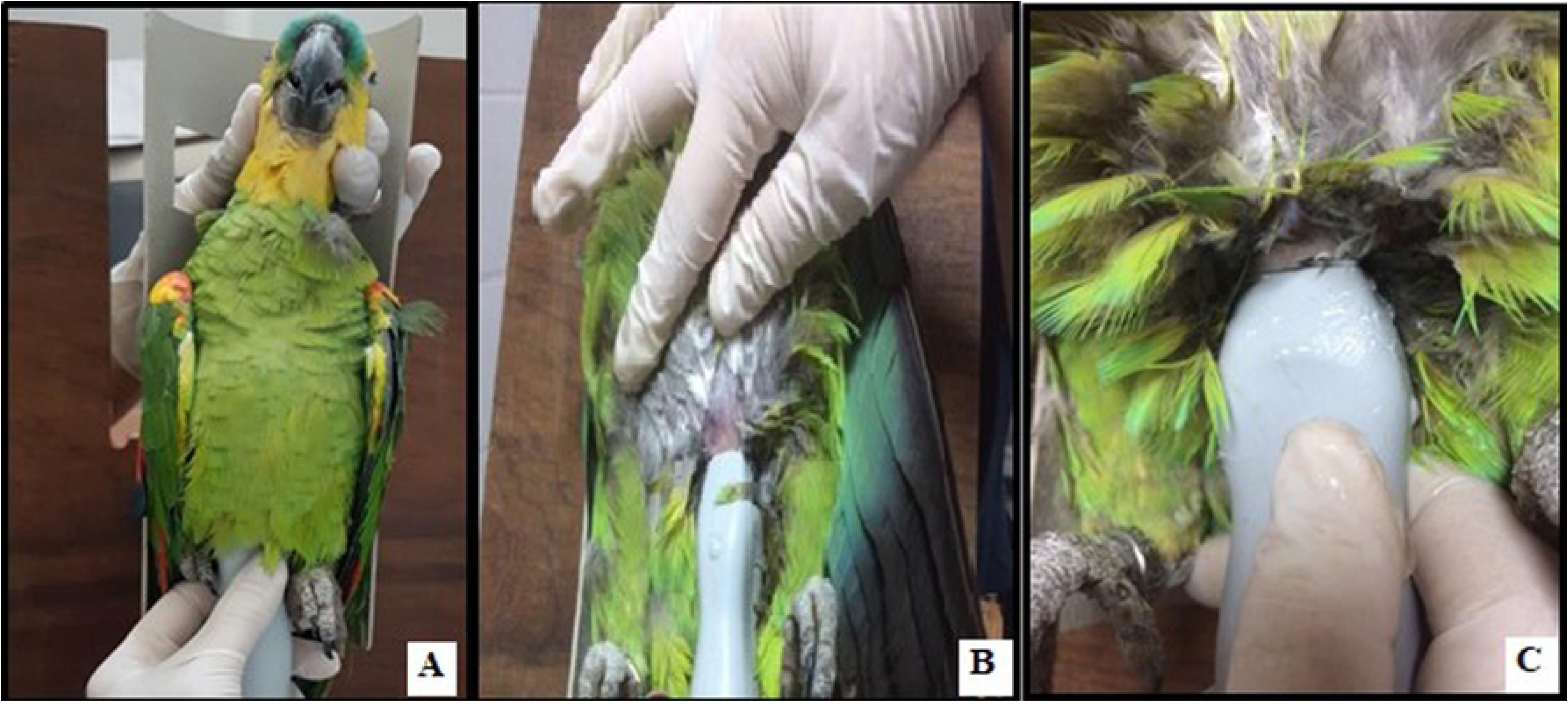
Positioning of bird and echocardiographic sections: A) Positioning of the bird for the echocardiographic examination with the aid of fixation device; B) Vertical echocardiographic section for visualization of two chambers; C) Horizontal echocardiographic section for visualization of all four chambers.

The ultrasound beam was directed craniodorsally, using the liver as an acoustic window. The heart was evaluated through two image planes [17]: the horizontal plane, used to assess the right and left atrial and ventricular chambers; and the vertical plane, used to assess the left atrial and ventricular chambers (Fig. 2A and 2B). The echocardiographic measurements were taken on the horizontal section, with the transducer positioned in such a way as to allow the visualization of the maximum expansion of the cardiac chambers, employing the inner border method at the end of diastole and systole.

**Fig 2:**
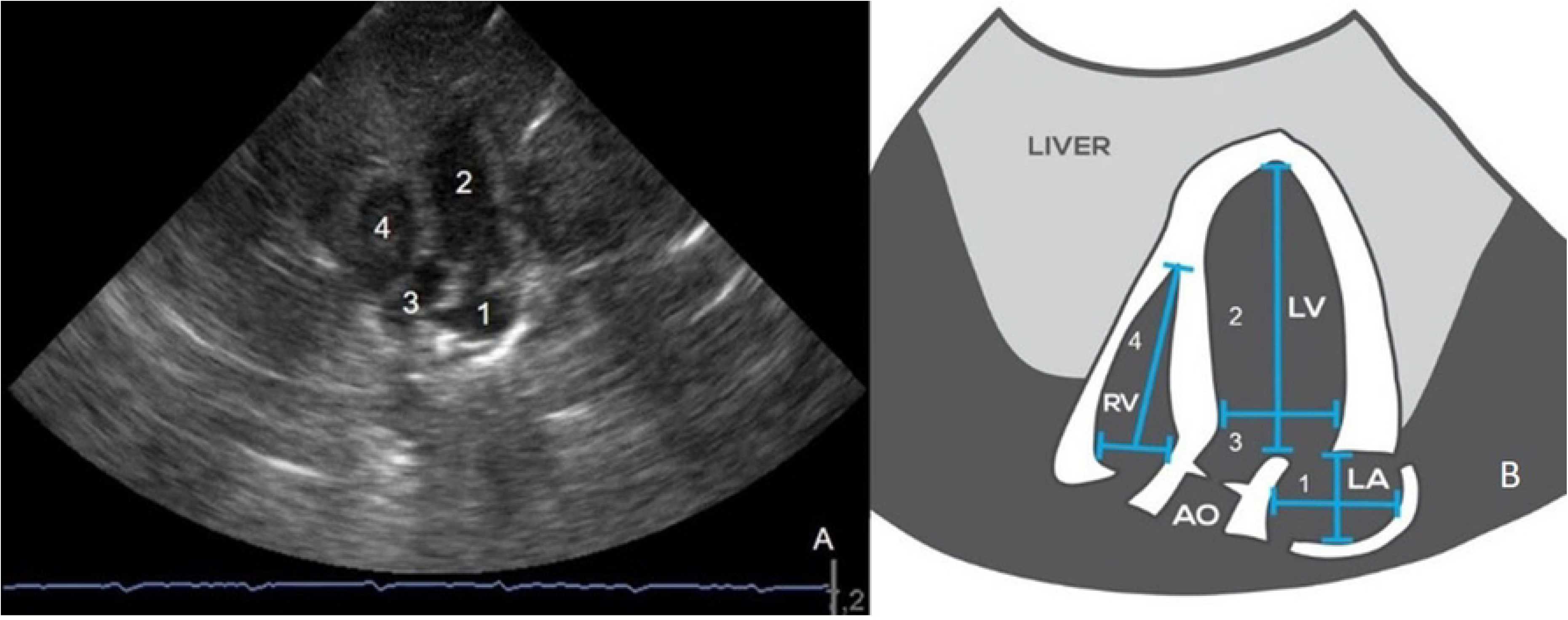
Echocardiographic horizontal plane and schematic echocardiographic representation. A) Echocardiographic section, horizontal plane showing: 1- left atrium (LA); 2- left ventricle (LV); 3- aorta (AO); 4- right ventricle (RV); B) Schematic echocardiographic representation, horizontal plane: 1- LA; 2- LV; 3- AO; 4- RV.

A subjective cardiac analysis was conducted, assessing the correlation between the ventricles, the thickness of the interventricular septum and ventricular contractibility. The echocardiographic measurements assessed were: length and width of the left and right ventricles during systole and diastole; length and width of the left atrium at the end of the systole. For the analysis of the systolic function, the fractional shortening (%) was determined for both ventricles through the following equations: [(DVW-SVW) / DVW] x 100, where DVW = diastolic ventricular width and SVW = systolic ventricular width; and [(DVL - SVL) / DVL] x 100, where DVL = diastolic ventricular length and SCV = systolic ventricular length. Three measurements were taken from different cardiac cycles and the mean value was calculated [15,17].

The computed tomography (CT) scan was conducted with a helicoidal SCT-7800 TC tomograph (Shimadzu; Kyoto, Japan). The acquisition protocol was 120kVp, 100mA (pitch of 1.5 with 1 mm increments and tube rotation time of 1 second), with a field of view (FoV) of 350×350 mm using a soft tissue window. The thickness of the sections was 3 mm [18]. The birds were positioned in dorsal decubitus with the pelvic limbs relaxed and the wings extended laterally. The scout view was used to locate the sections, demarcating the cranial and caudal limits for acquisition of the definitive axial images. Image acquisition was done in the rostrocaudal direction. The images were transferred to the software Voxar-3D (Barco; Edinburgh, Scotland) for multiplanar reconstruction (MPR) in the sagittal and dorsal planes, and then analyzed with the PACS system (Synapse, Fuji Medical System, Tokyo, Japan).

The radiographic examination was conducted using a direct digital radiography device (DR-F; GE Health Care Unit, Brazil), with the settings configured at 45 kV, 200mA and 5 mA with 250ms, broad focus, focal-film distance of 100 cm and collimation on the focal region for all projections. Image acquisition was conducted in the following projections: right lateral, with the wings overlaid and extended dorsally, and the legs extended caudally with the femoral heads overlaid; ventrodorsal, with the wings extended laterally and the legs extended caudally symmetrically. The images were analyzed with the PACS system (Synapse, Fuji Medical System, Tokyo, Japan).

The following measurements were evaluated in the radiography and CT scan: length of the heart (mm), from the base to the apex (lateral projection for radiography and sagittal section for the CT scan) (Fig 3A and 4A, respectively); width of the heart (mm) at the broadest part of the heart (ventrodorsal projection for radiography and dorsal section for the CT scan) (Fig 3B and 4B, respectively); width of the coelom (mm), from one rib to the other at the same level as the width of the heart for radiography (ventrodorsal projection) (Fig 3B) and at the largest width of the coelom for the CT scan (dorsal section) (Fig 4C); ratio between the width of the cardiac silhouette and the width of the coelom (%).

**Fig 3:**
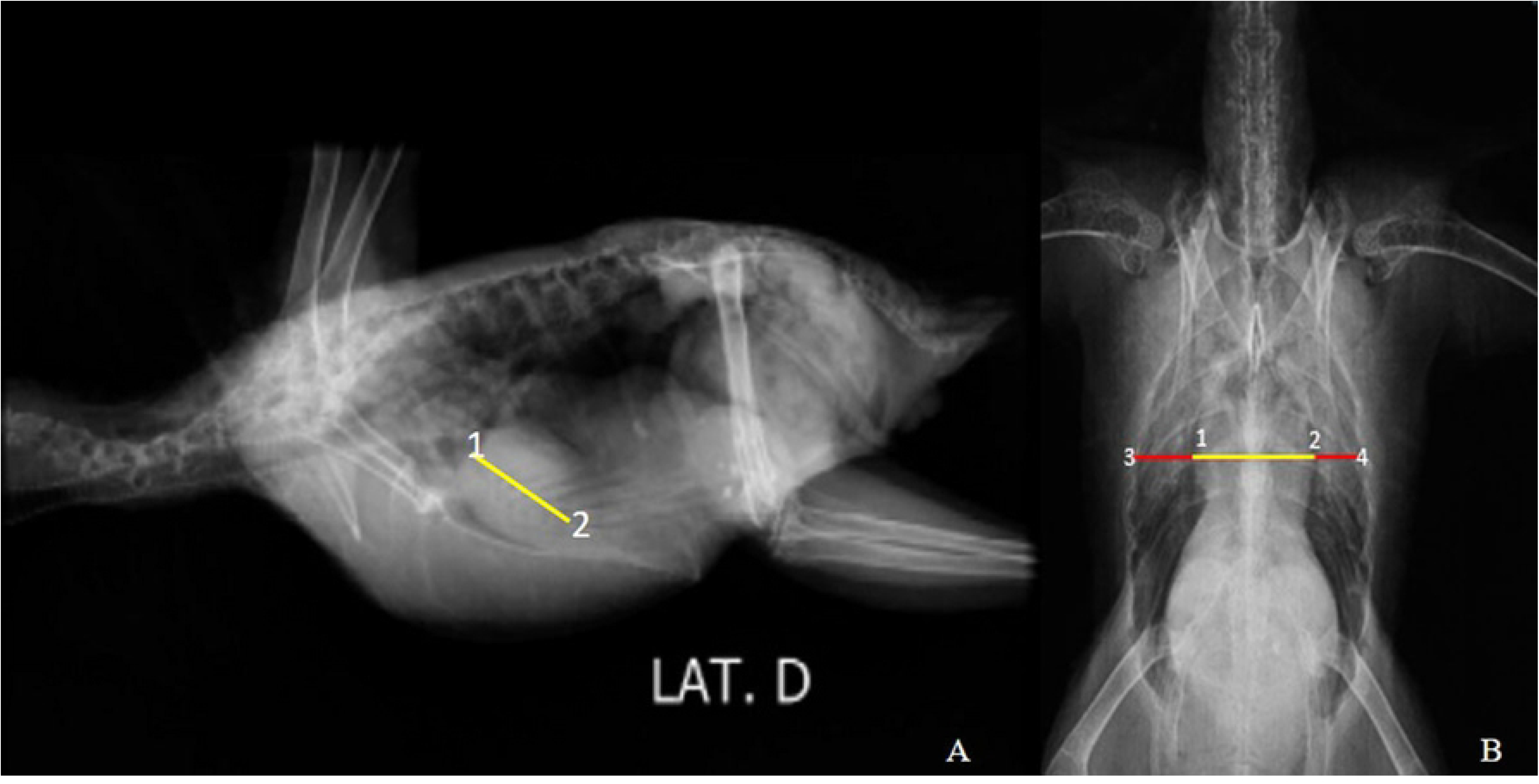
Radiographic image. A) length of the cardiac silhouette between points 1 and 2, right lateral projection; B) width of the cardiac silhouette between points 1 and 2; width of the coelom between points 3 and 4, ventrodorsal projection.

**Fig 4:**
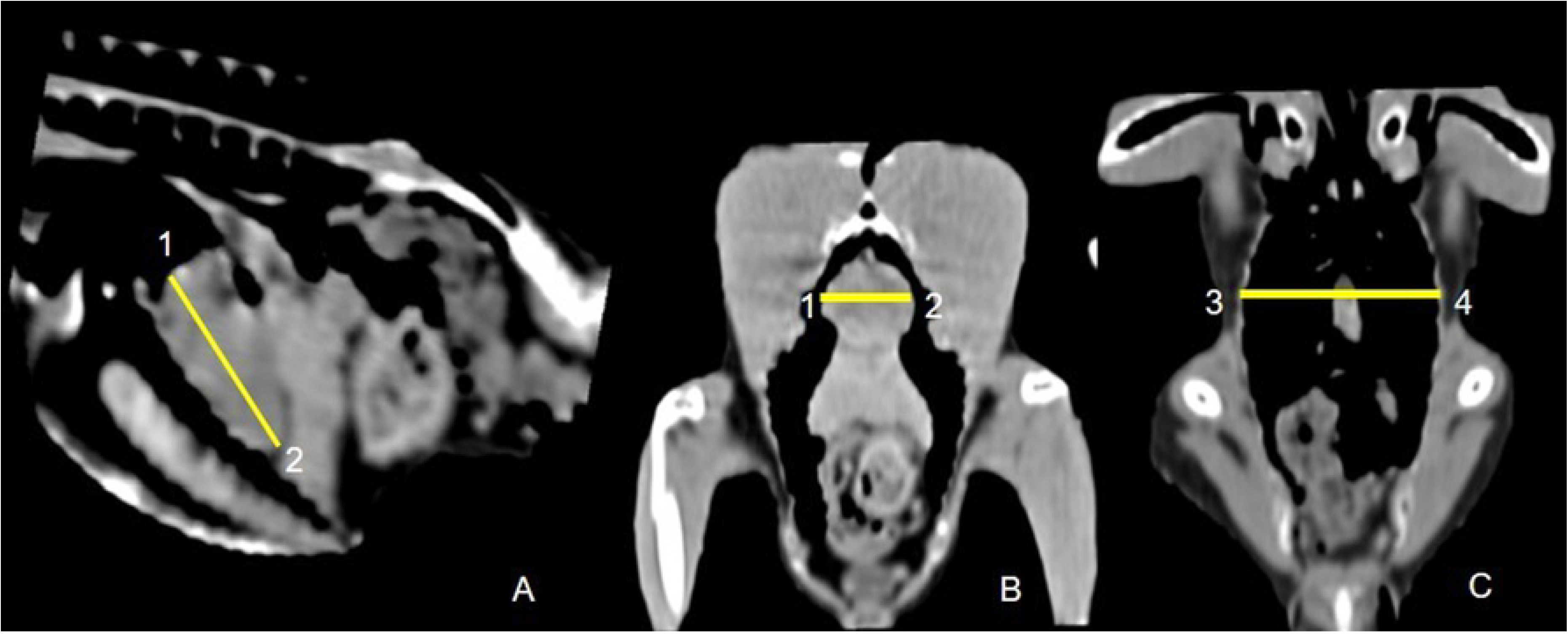
Tomographic image. A) length of the cardiac silhouette between points 1 and 2, sagittal projection; B) width of the cardiac silhouette between points 1 and 2, dorsal projection; C) width of the coelom between points 3 and 4, dorsal projection.

The comparison of the cardiac and biometric data between the three groups (Lean, Ideal and Obese) was performed with the one-way analysis of variance (One-way ANOVA) followed by Tukey-Kramer post-hoc test. Data normality was verified through the Kolmogorov and Smirnov test. For parameters without any significant differences between the groups, the data was grouped (Lean + Ideal + Obese). The significance level was defined at 5%.

## Results

The heart exams of 35 parrots (*A. aestiva*) were assessed in this study. Among the animals included in the study, 14 (40%) presented normal BCS (BCS = 3 and 4), 11 (31.4%) were lean (BCS = 1) and 10 (28.6%) were obese (BCS = 5). The average body weight (g) were 337.8 ± 23.3g, 404.2 ± 227.4 and 465.6 ± 32.6 for the lean, ideal and obese groups, respectively, and was significantly different between the three BCS. The average length (cm) of the birds in the lean, ideal and obese groups were 32.9 ± 3.3, 33.1 ± 3.0 and 34.8 ± 1.6 respectively, with no statistically significant difference between the groups. The anesthetic protocol adopted immobilized the animals for about 60 minutes, which was enough time to conduct all three examinations.

With the echocardiographic technique employed, it was possible to conduct all proposed measurements in the 35 parrots included in the study through the horizontal plane. There were no significant different between the groups in the measurements conducted on the horizontal section for the following parameters: length and width of the left ventricle in systole and diastole; length and width of the left atrium in systole and diastole; length of the right ventricle in systole; width of the right ventricle in systole and diastole. However, obese parrots presented shorter diastolic lengths in the right ventricle (6.9 ± 1.6) than lean parrots (9.1 ± 2.1) (p=0.03).

For measurements that did not present significant differences between the groups, the data was grouped and compared with the literature. Therefore, the values obtained in the echocardiogram through the horizontal section in this study were in line with those observed in psittacidae [17,19] and pigeons [20], except for the systolic and diastolic lengths of the left ventricle, which were lower than the ones reported in the literature (Table 1). It is important to note that these differences may be caused by the difference in species/subspecies studied and that the number of animals included in this study (n=35) is significant in comparison with the other studies on psittacidae (n=10) [17,19].

**Table 1:** Echocardiographic values obtained through the horizontal section in parrots and pigeons, both in the literature and in this study.

| **Authors** | **Beufrère et al.**<br>**(2016)** | **Pees et al.**<br>**(2004)** | **Krautwald-**<br>**Junghanns et**<br>**al. (1995)** | **Study Data** |
| --- | --- | --- | --- | --- |
| **Species** | ***Amazon***<br>***parrots (n=10)*** | ***Amazon***<br>***parrots (n=10)*** | ***Racing pigeons***<br>***(n=50)*** | ***Amazona***<br>***aestiva (n=35)*** |
| **LV** |  |  |  |  |
| Ls (mm) | 16.5 – 25.7 | 20.7 ± 1.5 | 17.9 ± 1 | 11.76±2.9 |
| Ld (mm) | 17.7 – 26.5 | 21.8 ± 1.9 | 20.1 ± 1.4 | 14.86±2.9 |
| Ws (mm) | 4.3 – 9.1 | 6.7 ± 1.1 | 5.2 ± 0.4 | 5.25±1.17 |
| Wd (mm) | 6.4 – 10.4 | 8.7 ± 1.2 | 7.4 ± 0.6 | 7.04±1.55 |
| FS (%) | 14.4-31.2 | 22.8 ± 4.2 | - | 25.31±13.28 |
| **RV** |  |  |  |  |
| Ls (mm) | 5.8 – 13 | 9.4 ± 1.8 | - | 7.1±2.57 |
| Ld (mm) | 7.7 – 12.9 | 10.3 ± 1.3 | 9.9 ± 0.8 | 7.36±1.81* |
| Ws (mm) | 1.7 – 4.5 | 3.1 ± 0.7 | 4.0 ± 0.5 | 2.37±0.77 |
| Wd (mm) | 2.6 – 7.8 | 5.2 ± 1.3 | - | 3.07±1.06 |
| FS (%) | 26.7 - 41.5 | 34.1 ± 3.7 | - | 25.71±14.37 |

LV: left ventricle; RV: right ventricle; Ls: length in systole; Ld: length in diastole; Ws: width in systole; Wd: width in diastole; FS: fractional shortening in the longitudinal view; \*n=24 (animals in the obese group were not included, which were lower than the animals in the lean and ideal groups).

It was possible to evaluate only the aortic flow in the parrots due to high heart rate that is characteristic to the species, image overlaying and hyperechogenicity. When assessing the echocardiographic variables related to flow, the aortic speed (p= 0.002) and the aortic pressure gradient (p=0.02) were lower in the lean group than in animals with ideal scores. The values observed for aortic speed (m/s) aortic pressure gradient (mmHg) were, respectively, 0.8 ± 0.1 and 2.6 ± 0.8 in the lean group; 1.0 ± 0.1 and 4.0 ± 1.1 in the ideal group; and 0.9 ± 0.1 and 3.3 ± 0.5 in the obese group.

The fractional shortening (FS) of the right ventricle (RVFS) and left ventricle (LVFS) did not present differences between the three groups studied. However, the values for FS of the transverse axis of the LV, FS of the longitudinal axis of the LV and FS of the longitudinal axis of the RV were considerably lower in obese parrots, although without any statistical significance (Table 2). The transverse RVFS did not present such reduction in the obese group.

**Table 2:** Fractional shortening of the right and left ventricles, according to length and width, in the different body condition score groups.

| **Fractional Shortening (Fs)** | **Lean** | **Ideal** | **Obese** | **p Value** |
| --- | --- | --- | --- | --- |
| FSLV Transverse (%) | $32.1 \pm 14.3$ | $37.5 \pm 14.8$ | $22.3 \pm 10.4$ | 0.07 |
| FSRV Transverse (%) | $34.8 \pm 14.3$ | $28.6 \pm 19.5$ | $29.9 \pm 13.4$ | 0.64 |
| FSLV Longitudinal (%) | $28.8 \pm 10.6$ | $28.1 \pm 15.8$ | $15.0 \pm 6.9$ | 0.057 |
| FSRV Longitudinal (%) | $29.5 \pm 15.8$ | $27.0 \pm 15.2$ | $17.4 \pm 7.7$ | 0.34 |

LV: left ventricle; RV: right ventricle.

The radiographic examination showed an overlap of the apex of the heart with the cranial portion of the liver silhouette, which presented a difficulty for taking the measurements of the heart in the lateral projection and made it impossible in the ventrodorsal projection due to a higher degree of overlap. Therefore, the length of the heart in the radiographic images was measured only in the lateral projection. On the other hand, the width of the heart and the width of the coelom were measured in the ventrodorsal projection.

No significant differences were observed when comparing the measurements between the groups, both in the radiography and in the tomography. Therefore, the values for all three groups were grouped together and are presented on Table 3.

**Table 3:**
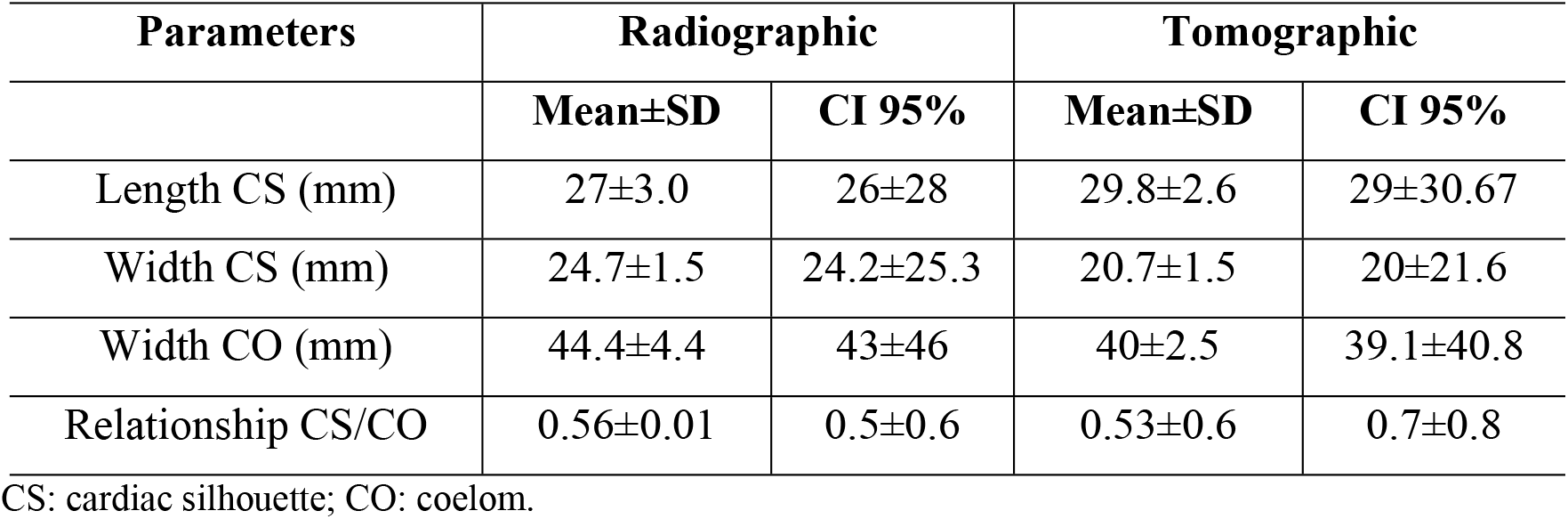
Mean, standard deviation (SD) and minimal and maximal confidence interval (CI 95%) for the radiographic and tomographic parameters for the grouped parrots (n=35).

| Parameters | Radiographic |  | Tomographic |  |
| --- | --- | --- | --- | --- |
|  | Mean±SD | CI 95% | Mean±SD | CI 95% |
| Length CS (mm) | 27±3.0 | 26±28 | 29.8±2.6 | 29±30.67 |
| Width CS (mm) | 24.7±1.5 | 24.2±25.3 | 20.7±1.5 | 20±21.6 |
| Width CO (mm) | 44.4±4.4 | 43±46 | 40±2.5 | 39.1±40.8 |
| Relationship CS/CO | 0.56±0.01 | 0.5±0.6 | 0.53±0.6 | 0.7±0.8 |

CS: cardiac silhouette; CO: coelom.

## Discussion

The anesthetic protocol employed in this study, with intravenous ketamine and midazolam, was efficient and immobilized the birds for a period of 60 to 80 minutes, which was enough time to conduct the echocardiographic, tomographic and radiographic examinations. The average duration of the examinations after sedation was 30 to 40 minutes for the echocardiogram, 20 minutes for the x-ray and 10 minutes for the tomography. The parrots presented a stable clinical profile, without any arrhythmias or alterations in the clinical parameters during anesthesia. The use of a sedation protocol that is safe and efficient is important to restrain the birds during clinical and diagnostic procedures because it reduces stress and facilitates handling [21].

Massone [22] reports that ketamine induces minimal events in the electrical conduction system of the heart, with the possibility of arrhythmias due to the increased sensitivity in the myocardium caused by the increased number of circulating catecholamines. According to the author, ketamine may lower myocardial contractility by interfering with the availability of intracellular calcium. However, the same anesthetic protocol was used in the three groups studied, which means the differences observed between groups may not be attributed to the anesthesia.

This anesthetic protocol allowed all birds to relax enough to be positioned correctly during the echocardiographic, radiographic and tomographic examinations. When conducting a radiographic examination it is extremely important to avoid overlapping the wings and pelvic limbs with the region of the heart [23]. The acetabula and humeroscapular joints should be overlapped in the lateral projection, while the sternum and the spine should be overlapped in the ventrodorsal projection, both of which were possible in this study. For birds, the optimal positions for radiographic examinations are the lateral and ventrodorsal positions [23].

Regarding the technique employed to acquire the echocardiographic images, physically restraining the animals with a wooden device allowed us to obtain the images with ease. Pees et al. [17] have suggested the use of a restraining device in psittacidae in order to position the birds without affecting the circulatory system and allow access to the contact area with the transducer. A similar device was employed in this study (Fig 2), which enabled the examination to be conducted more easily and allowed the transducer to be positioned horizontally and vertically.

The echocardiographic examination is one of the most important diagnostic tools in aviary cardiology [19]. According to Augusto [24], echocardiographic examinations in birds may be hampered by anatomical particularities, such as feathers, scales, air sacs and the thickness of the skin. The transducer was positioned at the ventromedial region, immediately posterior to the sternum, and it was possible to conduct the examination only by holding the feathers away from the device.

In the doppler echocardiogram, the size of the chambers and the contractility of the ventricles were assessed subjectively and through measurements. The morphology and function of the left and right atrioventricular valves, aortic valve and pulmonary valve presented alterations in the quality of the image, which, together with the elevated heart rate of psittacidae, did not allow the evaluation of the structures. In addition, the study was conducted with parrots of varying body scores. Fat deposits and the size of the animal may interfere with the positioning of the transducer, hindering the passage of the sound waves or the return of the echoes, which may lead to difficulties in obtaining the images [24]. In this line, we observed that obesity limited the accuracy of the images, but it was still possible to assess the proposed echocardiographic measurements in all parrots through the horizontal view.

In birds, the echocardiographic evaluation has been limited to the B-mode cardiac measurements because the M-mode, which allows the assessment of the fractional shortening and is routinely used in cats and dogs, cannot be reproduced in the species [25]. These limitations are caused primarily by the position of the heart in birds, withdrawn from the sternum and laterally surrounded by a system of air sacs, which does not provide an adequate window for the examination [25]. In addition, the ejection fraction was not assessed in the B-mode, as it would require a more precise image and a longer examination time, which would, in turn, require the animals to be restrained for longer periods.

Alternatively, transesophageal echocardiography improves the accuracy of the examination in M-mode [25] and in the detection of conditions such as endocarditis, intracardiac masses, thrombi and some heart diseases [26]. In birds, this modality improves the assessment of the structure and function of the heart, including the transverse transventricular view, which cannot be obtained with transcoelomic echocardiography. However, the technique should be employed with caution in psittacidae [25].

According to the echocardiographic data obtained in this study, we observed interference of the nutritional condition on the cardiac parameters. Obese parrots presented reduced right ventricle (RV) length in diastole and fractional shortening (FS). In obese humans, the function of the right ventricle has been associated with the dilation of the right atrium and ventricle, as well as increased thickness of the right ventricle free wall [27].

However, in this study the obese parrots presented lower dimensions in the diastolic length of the right ventricle than lean animals. It is important to note that the assessment of the right ventricle in other species is still the subject of many studies due to the difficulty in evaluating the chamber. In addition, the evaluation of the diastolic function is determined by parameters such as the E/A ratio, E-wave deceleration time (EDT) and isovolumetric relaxation time (IVRT), in addition to being influenced by the preload and presenting negative correlation with the heart rate (HR) [28]. Since these parameters were not assessed in the study, it is not possible to infer that the obese parrots suffered from diastolic dysfunction.

Obese parrots presented an apparent systolic dysfunction evidenced by the lower FS in the left ventricle, with values below [17] and also in line [19] with the standards for the species. Obesity in humans, cats and dogs is a volume-expansion disease with increased cardiac output, increased volume of plasmatic and extracellular fluids, increased cardiac chronotropism, systolic and diastolic dysfunction in the ventricles and increased arterial blood pressure [29].

The reduction of the fractional shortening (FS) in obese individuals in comparison with the other groups, although not statistically significant, clearly shows that the longitudinal FS of the left ventricle (LV) in the obese group (14.98 ± 6.88) was about half of the value observed in the lean (28.82 ± 10.59) and ideal score groups (28.12 ± 15.76) (p=0.057). These results are controversial, considering that in humans it has been recently described the preservation of the ejection fraction in obese patients with heart failure, known as the obesity paradox [30]. However, echocardiographic studies on birds are scarce and the values described in the literature for the LVFS (%) in *Amazona* sp. parrots are variable [17,19] (Table 1). It is also important to note that given the anatomical particularities in the heart and the differences in the standardized echocardiographic sections in birds, the comparison with humans and other mammals is not entirely valid. The reduced FS may be explained by factors such as increased afterload, reduced preload and low contractility, while increased FS may be explained by increased preload, reduced afterload and increased contractility [28].

Obesity increases the workload of the heart by increasing the total volume of blood and the cardiac debt [31]. The excess of fat tissue imposes an increase on the metabolic demand of the body, leading to a hyperdynamic circulation, structural changes (remodeling) in the right and left ventricles and, therefore, increased ventricular mass and cavity dilation. Obesity is associated with hypertrophy, dilation and diastolic dysfunction of the left ventricle and occasionally systolic dysfunction of the left ventricle [27].

In addition to the effects of obesity observed in this study, we also noted that weight loss and malnutrition in parrots also had an impact on the cardiac flow. We observed a reduction in aortic speed and pressure gradient in the Lean group, while the FS was equal or slightly higher than in the Ideal group. In dogs, inducting caloric and protein malnutrition promoted the reduction of the cardiac mass, reduction of the FS and reduction of the cardiac output. The output reduction was attributed to a decrease on the metabolic demands that happened in malnourished individuals [32]. However, there is a lack of studies assessing the influence of nutritional condition over the cardiac function of birds.

In this study, when the data obtained on the echocardiogram that did not show an influence of the body condition score were grouped (horizontal plane), the values were similar to those observed for psittacidae [17,19] and pigeons [20], except for the length of the left ventricle in systole and diastole, which were lower than the reference values (Table 1). This may be explained by the inconsistent size of the animals and by the different species and subspecies included in the studies.

In the radiographic examination, an ‘hourglass’ shape was observed at the area of the heat on the ventrodorsal projection (Fig 5). This is common in psittacidae [8] and is caused by the overlap of the heart and the liver due to the missing diaphragm in birds, which contributes to the formation of this ‘cardio-hepatic waist’ [33]. Alterations such as cardiomegaly, microcardia and microhepatia deform this silhouette and, therefore, the cardio-hepatic waist [8].

**Fig 5:**
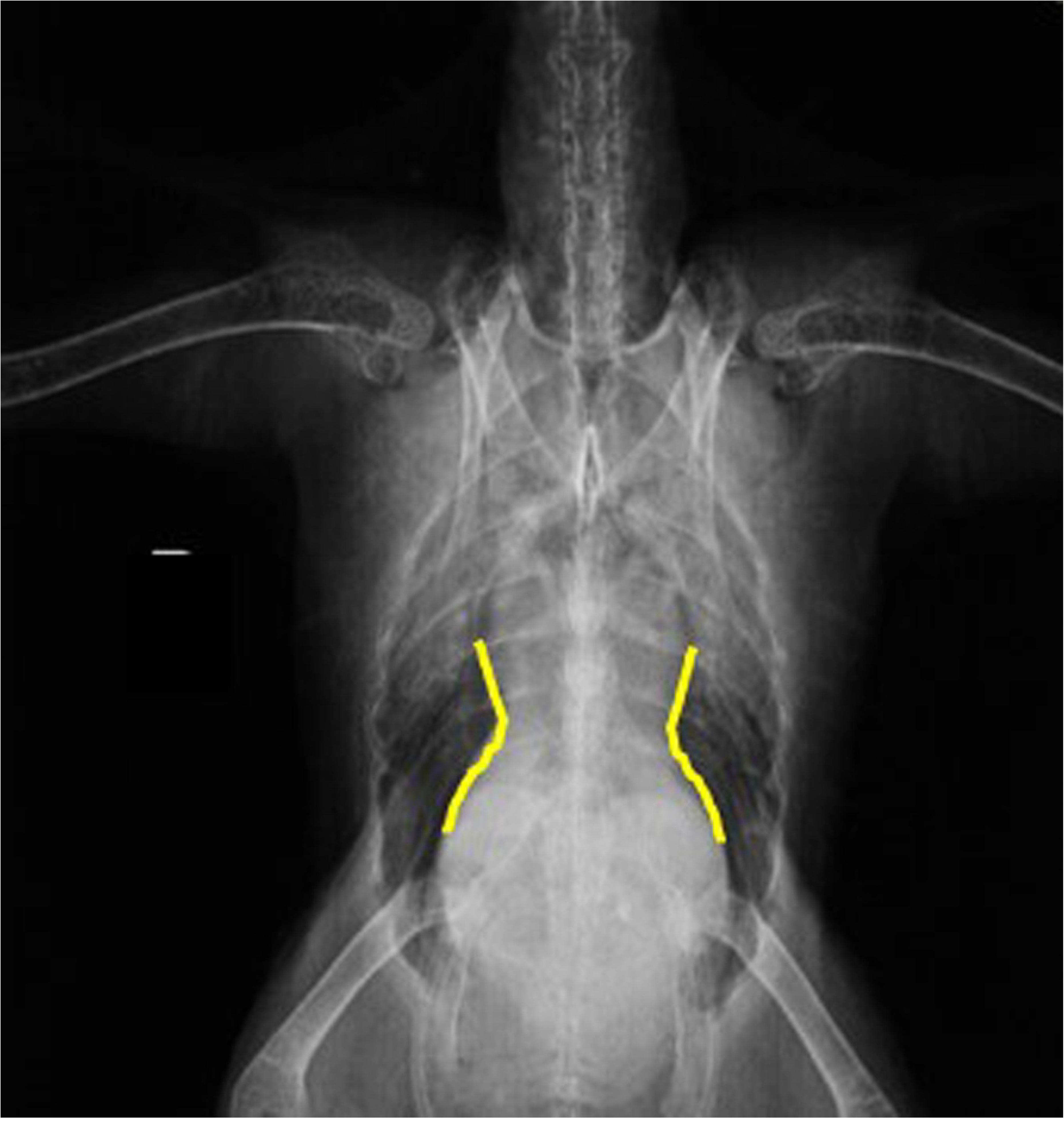
Radiographic image. Showing the hourglass shape created by the overlap between the apex of the heart and the cranial portion of the hepatic silhouette on the ventrodorsal position.

This overlap between heart and liver complicated the measurement of the length of the heart, which was ultimately measured only in the lateral projection and required adjusting the contrast of the images with the software. Different from radiography, computed tomography provides images in sections, without tissue overlaps since the images are obtained initially through axial sections and from these it is possible to create multiplanar reconstructions, such as the sagittal and dorsal planes [34]. Such reconstructions were used in this study to measure the length and width of the heart in the sagittal and dorsal sections, respectively. The measurements of the cardiac silhouette obtained through the CT scan were one of the pieces of information provided by this examination, which represents an additional tool for detecting possible alterations involving the heart.

The ratio between width of the heart and width of the thorax is considered normal for parrots (*A. aestiva*) at the range of 51 to 61% [35]. However, this study observed lower and higher values in all groups. Two lean animals (18.2%), one animal with ideal score (7.1%) and two obese animals (20%) presented results below the reference values, while one lean animal (9.1%), two animals with ideal scores (14.2%) and four obese animals (40%) presented results above the reference. Therefore, caution should be exercised when interpreting this ratio based on the data described in the literature.

The CT scan also measured this ration and the results did not diverge according to the body condition scores. When the animals were grouped together, the ratio varied between 43 and 62%, with a mean value of 53±6%. A review of the literature revealed no data regarding this ratio on tomography. While it was possible to assess the cardiac silhouette through the CT scan, the atrial and ventricular chambers were not visible. According to VELADIANO et al. [3], it was possible to view all four chambers in macaws, but intravenous contrast was used, which did not occur in the present study. When comparing the width of the cardiac silhouette, the width of the coelom and the ratio between the two in the radiographic and tomographic images, we noted that the values were higher in the radiography. However, this difference was expected considering that the image formation methods are different in both examinations, with radiography images being bidimensional and CT images tridimensional [36]

One of the limitations of this study was the non-use of iodized contrasts during the tomography of the chart. In humans, cardiac CT scans involve the use ionizing radiation and iodized contrasts, representing an important non-invasive method for the diagnosis of atherosclerotic disease [37]. Psittacidae present a high incidence of atherosclerosis [38], a common cardiac disease that is often associated with sudden death to which birds appear to be more susceptible than mammals [39]. This fact helps illustrate the importance of contrasted tomography for the species.

## Conclusions

Therefore, we may conclude that changes in the nutritional condition of blue-fronted Amazon parrots (*A. aestiva*) may lead to cardiovascular dysfunctions detectable only through echocardiographic examination, which represents an important diagnostic tool for the species. Although the CT scan allowed a better assessment of the structures of the cardiovascular system, the radiographic examination still should be considered the standard screening test to identify cardiac alterations such as increased or reduced dimensions. Describing the techniques used and the measurements obtained in this study may contribute towards further research in the area.

## Acknowledgements

The authors thank everyone involved in this study and specifically the Wildlife Medicine and Research Center (*CEMPAS*, *Centro de Medicina e Pesquisa em Animais Selvagens*) and School of Veterinary Medicine and Animal Science, UNESP, Botucatu, Brazil.

## Author Contributions

**Conceptualization:** Gisele Junqueira dos Santos, Amanda Sarita Cruz Aleixo, Jeana Pereira da Silva; Alícia Giolo Hippólito; Bárbara Sardela Ferro,; Elton Luis Ritir Oliveira; Priscylla Tatiana Chalfun Guimarães Okamoto and Alessandra Melchert

**Data curation:** Gisele Junqueira dos Santos, Amanda Sarita Cruz Aleixo, Jeana Pereira da Silva; Alícia Giolo Hippólito; Bárbara Sardela Ferro,; Elton Luis Ritir Oliveira; Priscylla Tatiana Chalfun Guimarães Okamoto and Alessandra Melchert

**Formal analysis:** Gisele Junqueira dos Santos; Amanda Sarita Cruz Aleixo; Jeana Pereira da Silva; Sheila Canavese Rahal and Alessandra Melchert

**Investigation:** Gisele Junqueira dos Santos and Alessandra Melchert

**Methodology:** Amanda Sarita Cruz Aleixo; Gisele Junqueira dos Santos; Jeana Pereira da Silva; Alícia Giolo Hippólito; Bárbara Sardela Ferro and Alessandra Melchert

**Project administration:** Maria Lucia Gomes Lourenço and Alessandra Melchert

**Supervision:** Alessandra Melchert

**Writing-original draft:** Alessandra Melchert and Gisele Junqueira dos Santos

**Writing-review & editing:** Alessandra Melchert, Maria Lucia Gomes Lourenço and Vania Maria de Vasconcelos Machado

